# Time-based binding as a solution to and a limitation for flexible cognition

**DOI:** 10.1101/2021.10.09.463761

**Authors:** Mehdi Senoussi, Pieter Verbeke, Tom Verguts

**Author notes:** MS and PV contributed equally to this work as first co-authors. **Corresponding author:** Dr. Mehdi Senoussi, Department of Experimental Psychology, Ghent University, Henri Dunantlaan 2, 9000, Ghent, Belgium.

## Abstract

Why can’t we keep as many items as we want in working memory? It has long been debated whether this resource limitation is a bug (a downside of our fallible biological system) or instead a feature (an optimal response to a computational problem). We propose that the resource limitation is a consequence of a useful feature. Specifically, we propose that flexible cognition requires time-based binding, and time-based binding necessarily limits the number of (bound) memoranda that can be stored simultaneously. Time-based binding is most naturally instantiated via neural oscillations, for which there exists ample experimental evidence. We report simulations that illustrate this theory and that relate it to empirical data. We also compare the theory to several other (feature and bug) resource theories.

## Introduction

The existence of resource constraints on cognition is undebated: Just consider listening to a long list of grocery items to be fetched, and heading off to the supermarket without a piece of paper (or smartphone) to support your memory. What is debated, however, is the *nature* of these resource constraints. Of course, resource is a broad term that has been applied throughout psychology and neuroscience (e.g., Barlow (1961)). However, we will restrict our attention to theories with immediate implications for working memory (e.g., as in the supermarket example). With this delineation out of the way, we note that a long research tradition has empirically investigated the nature of resource constraints (Bays & Husain, 2008; Cowan, 2001; Miller, 1956; Oberauer & Lin, 2017) by positing a limited quantity of some sort, and then deriving predictions (perhaps supported by a formal model) with respect to behavioral data in the working memory domain. This is the ‘bug’ approach mentioned in the abstract. However, in line with David Marr and the ‘feature’ approach, we first consider what a computational perspective would stipulate for flexible cognition(Holroyd & Verguts, 2021). To be clear from the start, “computational” is often used as in “instantiated in a formal model”; this is not what we mean here. By computational, we refer to the computations that are required in tasks relying on flexible cognition (such as getting one’s groceries, in the upcoming example). Our detour into flexible cognition lays the groundwork for our main thesis: The resource constraint is a consequence of the computational requirements to implement flexible cognition. Then, we consider the implementational perspective, and present some simulations to illustrate our theory, based on a recent oscillatory model of working memory (Pina et al., 2018). Finally, we relate our theory to other (similar and different) proposals in the General Discussion section.

### Role-filler binding

Cognition requires the flexible binding and unbinding of two or more elements. For example, an experimenter may instruct a subject to detect the red squares in a stream of stimuli, but ignore the blue squares and red triangles (Treisman & Gelade, 1980). More mundanely, a mother may ask her son to go to the store to buy a pack of gluten free pasta and 1kg of apples. If he comes home with 1kg of regular pasta and a pack of gluten free apples, he is likely to be sent back. As another example, acting appropriately in a restaurant requires binding the waiter role to the person running around with the drinks; this binding allows one to know how, what, and when to order. In a sense, cognitive life is built on binding.

A particularly important type of binding is that between roles to be filled and fillers of those roles (*role-filler binding*; Hummel (2011)). For example, suppose one wants to memorize that the fruit aisle is on the left of the dry food store in the supermarket. The roles are here “Left” and “Right”; the fillers are “fruit aisle” and “dry food department”; and the relevant role-filler bindings are (Left, fruit aisle) and (Right, dry food department). As an aside, these roles can be implemented via different types of representational codes, including verbal or spatial (Gevers et al., 2010; van Dijck & Fias, 2011); we currently remain agnostic about their nature. Consider as another example of role-filler binding, syntactic constructions such as the Subject – Verb – Object (SVO) type sentence. For example, in a sentence like “Tom buys pasta”, the relevant role-filler bindings are (Subject, Tom), (Verb, buys), and (Object, pasta). Other syntactic constructions are possible to represent this information (e.g, (Buyer, Tom), (Object-bought, pasta)), but the syntactic structure doesn’t matter for our argument, and we will stick to SVO constructions to explain our argument. We will discuss a few constraints on role-filler bindings in cognition, and how these constraints impose processing bounds on cognition.

Some sentences (such as “I love you”) occur sufficiently frequently to be stored as a separate chunk in memory, independent from other information. There is indeed evidence that such (high-frequency) chunks are important in language (McCauley & Christiansen, 2014), and perhaps in cognition more generally. However, chunking is not a realistic possibility for coding SVO sentences in general. For example, if there are *N* possible fillers (Tom, buy, book, …) and three possible roles (Subject, Verb, Object), a systematic chunking approach confronts a combinatorial problem, as it would require storage of 3*N*^2^ chunks of knowledge. More importantly, because information is stored independently in memory, a chunking approach does not easily lend itself to generalization (Marcus, 2001, 2018). If one learns something about books, flexible cognition requires that this novel information generalizes to the statement “Tom buys a book” (Fodor & Pylyshyn, 1988). For example, even a rudimentary knowledge about books is enough to conclude that buying a book entails a very different process than buying a house. But if the proposition that “Tom buys a book” is stored as a separate chunk in memory, such generalization between propositions is not possible.

The solution to this generalization problem involves compositionality (Fodor & Pylyshyn, 1988): Storing all components (or building blocks, here, roles and fillers) separately, in such a way that they can later enter into novel relations with other components. Applied to roles and fillers, this principle is also called *role-filler independence* (Hummel, 2011). Indeed, if one stores “book” information separately, the concept can later be independently enriched; and the novel information (e.g., that a book can be bought in bookstores, without the hassle and administration involved in buying a house) can thus be applied to instances like “Tom buys a house”.

With role-filler independence, the memory requirements are much lighter than in a chunking approach. Consider Figure 1: Here, *N* fillers and 3 roles are represented, with a much lighter memory requirement of just 3 + *N* elements. Any specific sentence (“Tom buys a book”) involves a combination of the corresponding roles and fillers.

**Figure 1.**
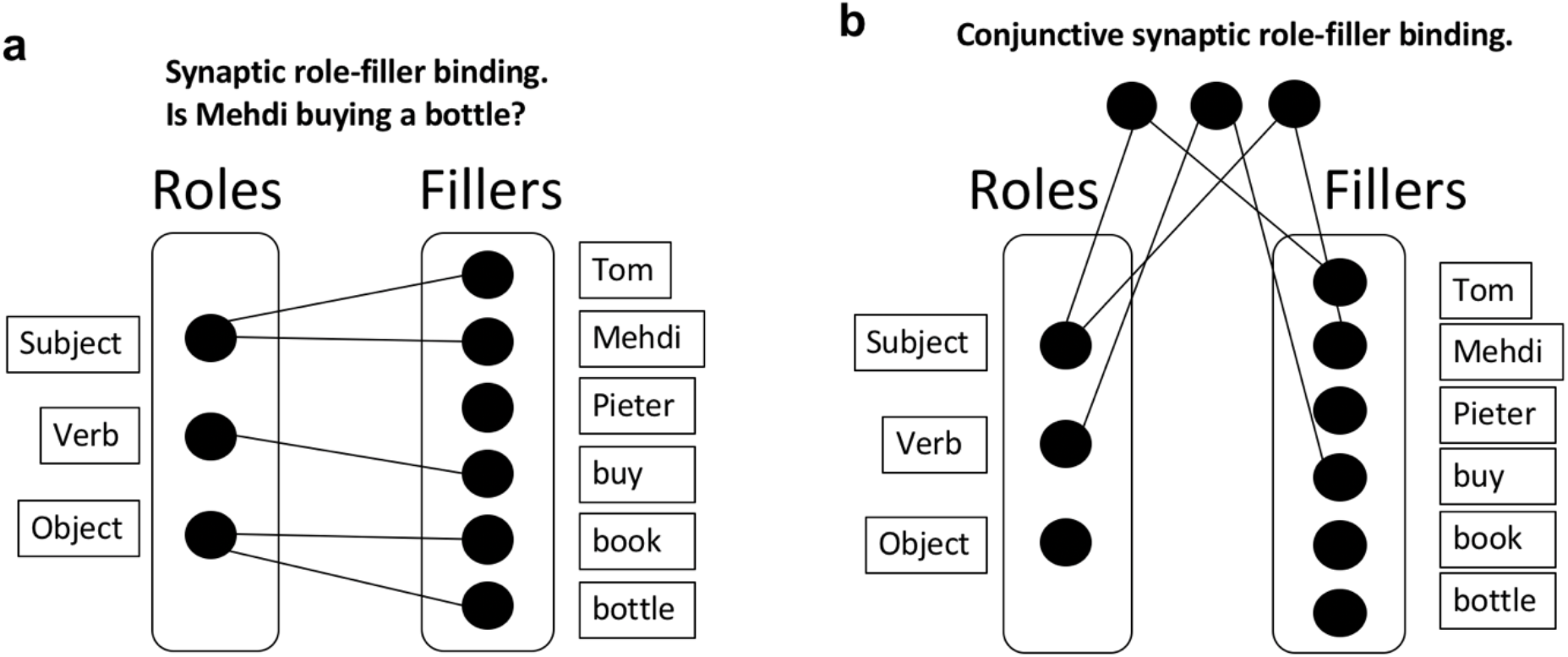
Synaptic binding. **a.** Synaptic role-filler binding. This approach allows to bind roles and fillers but has an important downside: after the presentation of multiple sentences, the roles and fillers are soon saturated, connecting all roles with all fillers. If synapses are not immediately removed, this approach will lead to interference; which we refer to as the unbinding problem. **b.** Conjunctive synaptic role-filler binding. An alternative solution is to implement a set of “gates” that would, for instance, activate specific roles and fillers. Once activated, a gate would activate the corresponding (role or filler) elements. This approach has the same unbinding problem, and additionally presents a combinatorial problem: 3N^2^ gates must be created to store all possible bindings.

If roles and fillers are stored independently, a next crucial property for flexible cognition is *dynamic role-filler binding* (Hummel, 2011). Specifically, it must be possible to rapidly bind and unbind roles and fillers in order to understand complex events in the real world. Consider hearing the story that Tom buys a book, then that Mehdi buys a bottle, and finally that Tom gives his book to Mehdi. In order to understand the three events and their logical relations, and in order to answer questions about the situation (Who currently owns two items?), it is important to initially bind Tom to Subject and book to Object; and then bind Mehdi to Subject (and unbind Tom and Subject) and bind bottle and Object (and unbind book and Object); and so on.

### Synaptic binding

It then remains to be explained how dynamic role-filler bindings are formed. One approach is to construct a synaptic connection between each (role, filler) pair for each sentence that is currently of relevance. We will call this a *synaptic binding* approach; presumably, a configuration of synapses stores the relevant information. Note that the term “synapse” can be interpreted either literally in the biological sense, or more metaphorically; the only functional requirement of a synapse for our purposes is that two memory elements are connected. It is the approach applied, for example, in neural network training algorithms (e.g., backpropagation; Bishop (1995)). Whereas originally thought to contribute mainly to long-term memory, recent work suggests that synaptic binding also supports working memory (the synaptic model of working memory; Stokes (2015)). However, this synaptic binding approach has its downsides. In particular, if synapses are not immediately removed after a sentence, interference will quickly occur. Consider again first representing that Tom buys a book, then that Mehdi buys a bottle, perhaps followed by some other purchases and exchanges of goods. In this case, roles and fillers will soon be saturated, connecting all roles with all fillers, and thus abolishing any meaning (see Figure 1a as an example). We will call this the unbinding problem of synaptic binding. Computationally, the problem manifests itself in catastrophic interference between partially overlapping tasks, which is a huge problem in neural networks (French, 1999), with several solutions being proposed to overcome it (Kirkpatrick et al., 2017; McClelland et al., 1995; Verbeke & Verguts, 2019). Also biologically, it’s not clear that the construction and destruction of biological synapses can occur at the time scale required for cognitive processing (Caporale & Dan, 2008; Kasai et al., 2003).

One could argue that the problem in the previous scenario derives from the direct synapses between roles and fillers. Thus, an alternative synaptic binding solution could be to implement a set of “gates” that filter out or activate specific roles and fillers. For example, there could be one gate for Tom, one for Mehdi, another for Subject, and so on. When the corresponding gates are activated, they in turn activate their corresponding (role or filler) elements. This approach would obviate the requirement of direct bindings between roles and fillers. However, in this approach, suppose each gate is selective for a specific role or filler; then appropriate (role, filler) pairs cannot be kept apart. Consider for example representing that Tom buys a book and Mehdi buys a bottle; in such a system, the interpreter of the system has no way to know whether the activated book belongs with Tom or with Mehdi. To solve this problem, one could suppose, instead, that there is a separate gate for each (role, filler) pair (Figure 1b). This approach could solve the problem of disambiguating different meanings, but here the combinatorial problem (3*N*^2^ gates must be created) and the unbinding problems appear again. We conclude that a pure synaptic binding approach is likely insufficient to implement dynamic role-filler bindings at the time scale required in systematic cognition.

### Time-based binding

Instead of synapses, one could consider using the time dimension to bind roles and fillers; we will call this a *time-based binding* approach. In particular, suppose that at time *t*, the role-filler pair (Subject, Tom) is active. However, a single time point doesn’t leave enough time for processing; moreover, the system doesn’t necessarily know when exactly the information will be of use in further task processing. It is thus useful to repeat the information for some period of time. Let’s suppose that the binding is repeated at intervals of length *d*. Hence, at all times *A*_1_ = {*t*, *t* + *d*, *t* + 2*d*, …} both elements of the (role, filler) pair (Subject, Tom) are active. Note that Subject and Tom are indeed joined by time only; there is no synaptic connection between them.

Besides representing (Subject, Tom), we also need to represent the pair (Verb, Buy). However, if the pair (Verb, Buy) were active at the same time as (Subject, Tom) (say, at time *t*) we run the risk of interference, as explained in the previous paragraph. We must thus represent it at some other time, say *t* + *e*. Just like for (Subject, Tom), we repeat the (Verb, Buy) pair at the same distance *d* so the two pairs maintain their temporal separation. Thus, we conclude that at times *A*_2_ = {*t* + *e*, *t* + *d* + *e*, *t* + 2*d* + *e*, ..} the pair (Verb, Buy) is active1. With a similar logic, at times *A*_3_ = {*t* + 2*e*, *t* + *d* + 2*e*, *t* + 2*d* + 2*e*} the pair (Object, Book) will be active.

### Synaptic learning on time-based bindings

It is well known that a neural network training rule (such as backpropagation) can learn complex tasks via synaptic binding, especially if it has available appropriate (here, compositional) representations of the input space. We postulate that this allows an efficient combining of the synaptic and time-based approaches. Specifically, once a time-based binding system as sketched above is constructed, a synaptic learning rule operating on its representations can subsequently learn various tasks. For example, in the book-buying context, the training rule could learn to answer questions such as “Who bought a book?”; “Who owns a book?”; and so on. Or in an experiment context, relevant mappings to be learned could be “Press the f key if you see a red square, the j key if you see a blue circle, and nothing otherwise”.

Importantly, such a representational system with independent and dynamic role-filler pairs, allows for generalization. For example, if novel information is learned about, say, Tom, this novel information can be attached (by the learning rule) to the representation of Tom, and thus be immediately generalized to other contexts in which Tom may appear.

Moreover, during both learning and performance, it’s very easy to delete old, no longer relevant information without leaving any trace to be erased: No synapses were created for binding, so none need be erased. It’s straightforward to represent the fact that Tom buys a book, followed by the fact that Mehdi buys a bottle. Finally, it’s relatively easy to construct new (role, filler) pairs via synchronizing “bursts” (Verguts (2017); and see Discussion).

In summary, we propose that synaptic and time-based binding ideally complement each other for the purpose of flexible cognition. Time-based binding allows quickly constructing and destructing connections. In contrast, synaptic binding allows application of very powerful learning rules. In this way, advantages of both synaptic and time-based binding are exploited, and their respective disadvantages are mitigated.

### A resource bound to time-based binding

Despite its several advantages, there is a constraint to the time-based approach. Specifically, this system of representing information will only work if the elements of sets *A*_1_, *A*_2_, and *A*_3_ (i.e. the timings of the different (role, filler) pairs) remain sufficiently separate (where “sufficient” depends on the level of precision required to robustly pass a message to a downstream neural area). Hence, such a system of representations can efficiently represent information (via dynamic (role, filler) bindings), and forget old, no longer relevant information. But it has an inherent constraint: It can only represent a limited number of elements at the same time.

Can we characterize this constraint more precisely? Note first that the first item of each set should be smaller than the second element of all sets (i.e. all distinct pairs/elements are activated once before any of them gets reactivated); otherwise, there is an ambiguity which set an element belongs to. In other words, we require that *t* + *ne* < *t* + *d*, that is, *n* < *d* / *e*. Storage capacity *n* has thus an upper bound *d*/*e*, determined by the period (*d*) of each set, and the time (*e*) between elements. This bound cannot be made arbitrarily high: If *d* is too high, the time between different elements ((role, filler) bindings) is too long, and the items cannot be simultaneously processed by a downstream neural area that must interpret the bindings. Imagine having to remember a grocery list with several minutes between the different items. Similarly, if *e* is too small, the separate elements cannot be disentangled from each other, either because of noise or because of the time scale of the downstream neural area. We propose that these factors together impose a bound on how many novel items agents can store simultaneously.

Note that the argument is purely computational: Any agent (biological or artificial) who is confronted with a task with the described requirements (simultaneous but systematic storing of possibly rapidly changing facts, using a single representational space), should use time-based binding; and as a result, he or she is subject to the constraints. The argument also clarifies when the bound applies and when it does not. It is perfectly possible to store thousands of facts via synaptic binding, as long as they do not require on-the-fly constructions and destructions of conjunctions of information. In other words, the resource bound applies to (non-synaptic) working memory, not to long-term memory.

## Simulation

We next consider how a time-based binding system may be neurally implemented. To construct such a system for representing novel, on-the-fly constructions, one needs a periodic or oscillatory function (i.e., *f*(*X* + *c*) = *f*(*X*) for some *c* and for all *X*)). The simplest choice is perhaps a sinusoidal (sine or cosine) function, but this is not necessary.

Consistently, a long research tradition has suggested an important role of oscillatory functions (oscillations in short) for cognition in general, and for binding elements in memory in particular (Gray & Singer, 1989). It is well known that coupled excitatory and inhibitory neurons can easily be employed to generate oscillations (Gielen et al., 2010; Wilson & Cowan, 1972). The coupling parameters (synaptic weights) between excitatory and inhibitory neurons determine the characteristics of the oscillations, such as their phase, amplitude and frequency.

To illustrate our argument, we used a recently proposed architecture of binding through oscillations to model working memory (Pina et al., 2018). This oscillatory neural network can bind and maintain elementary features (each represented by one node of the network) over time, while keeping different bindings apart. Each node of the network is composed of three components (a neural triplet, see Figure 2a). The three components are fast-excitatory (*u,* emulating AMPA synapses), slow-excitatory (*n*, emulating NMDA synapses), and inhibitory (*v*, emulating GABA synapses), respectively. The fast-excitatory - inhibitory pair constitutes a Wilson-Cowan type system (Wilson & Cowan, 1972). This pair exhibits limit cycle behavior (i.e. oscillations) and, as stated above, the characteristics of these oscillations (e.g. amplitude, frequency) can be controlled by changing the coupling weights between these components. Additionally, the slow excitatory component provides excitatory input to the inhibitory and fast excitatory components, thereby allowing bistability of the neural triplet (Lisman et al., 1998): an inactive state with low amplitude fluctuations, and an active state with persistent high amplitude oscillations. The left part of Figure 2a shows the connectivity between each component of a neural triplet; see Supplementary Information for a full description of the differential equations defining each component’s activity, and the value of each parameter including the weights between nodes of the network. We kept all parameter values equal to the main simulations in Pina et al. (2018), and only varied the inhibitory component’s time scale (*τ*_*v*_, Equation 2 in Supplementary Information). This parameter defines the speed at which the inhibitory component’s activity is updated. Varying *τ*_*v*_ allows to manipulate the oscillatory frequency of the nodes’ active state. Moreover, in this model, nodes form a network (upper-right part of Figure 2a) in which all fast-excitatory components excite each other, and all inhibitory components inhibit each other (see bottom-right part of Figure 2a). This connectivity allows these nodes to form a network that exhibits binding by phase; that is, when the peak of two nodes coincide, within a certain temporal range that we will call a ‘binding window’, they attract each other and align their peaks, forming a bound state. This network is further also characterized by competition between active nodes (or between groups of nodes that are bound together); i.e., when the peak of two nodes are separated by an interval outside of the binding window range, they will repel each other and remain active in an out-of-phase state. Due to the intrinsic attracting and repelling dynamics of this architecture, it can thus bind and maintain information to form distinct memories (a single active node, or bindings between nodes), while avoiding mixing them, through time-based binding. Each memory consists of one or multiple bound elementary features, each represented by a node. In line with the theory postulated above, a memory is activated only periodically (see Figure 2b).

**Figure 2.**
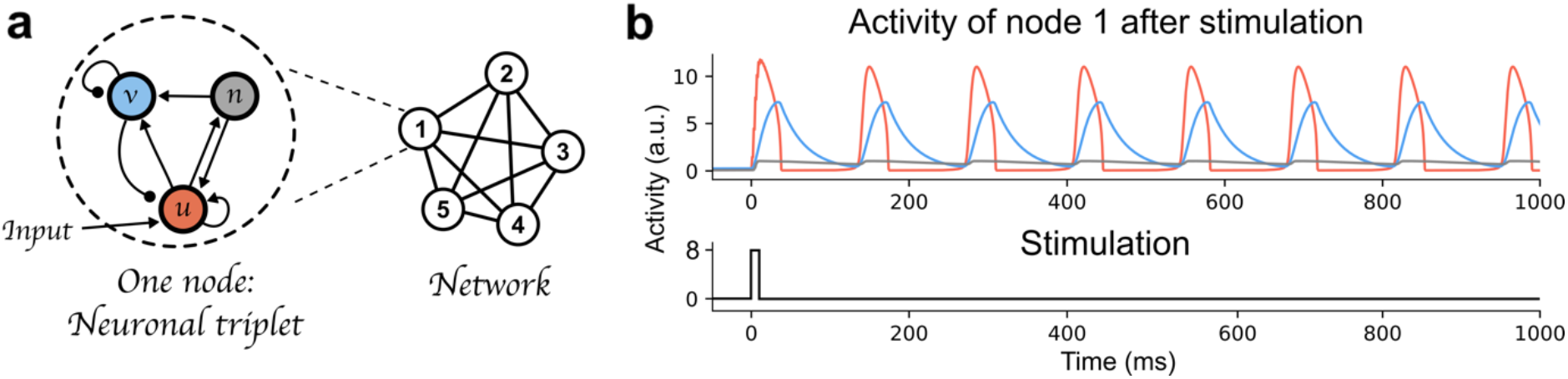
Time-based binding network. **a.** Each node of the network is a “neural triplet” composed of three components: a fast excitatory (u) neuron, an inhibitory (v) neuron, and a slow excitatory (n) neuron. Stimulation to a node affects unit u (as represented by the “Input” arrow). This architecture allows each node to start oscillating thanks to the excitatory-inhibitory pair of neurons (u and v, respectively), and maintain this oscillation through time (i.e. very slow decay) due to the slow excitatory neuron (n). When interconnected through excitatory-to-excitatory, and inhibitory-to-inhibitory connections, these nodes form a network that exhibits binding by phase and competition between active nodes or bound group of nodes. Synaptic weights are represented as lines between components or nodes: arrow ended lines represent excitatory connections, circle-ended lines represent inhibitory connections. **b.** Top: Example of node 1 being activated by an input stimulation. The red curve represents the activity of the fast-excitatory neuron (u), the blue curve represents the activity of the inhibitory neuron (v), and the gray curve represents the activity of the slow-excitatory neuron (n). Bottom: stimulation time course.

The ability to concurrently store multiple items of information in this manner, relies on two important features. First, the elements of each item must be bound together. For instance, nodes representing the role “Subject” and the filler “Tom” are in synchrony. Second, to concurrently maintain multiple memories (e.g., Subject-Tom and Verb-Buy), the two nodes representing the Subject-Tom pair must remain out of synchrony with those representing the Verb-Buy pair (see units 1-2 and 3-4 in Figure 3). This mechanism entails that the number of distinct memories that can be maintained simultaneously without interference, is limited by the frequency of the oscillation. In doing so, this model exemplifies how capacity limits emerge as a property of a system using oscillatory mechanisms for binding, i.e. the time interval between two peaks of activated nodes (or groups of nodes).

**Figure 3.**
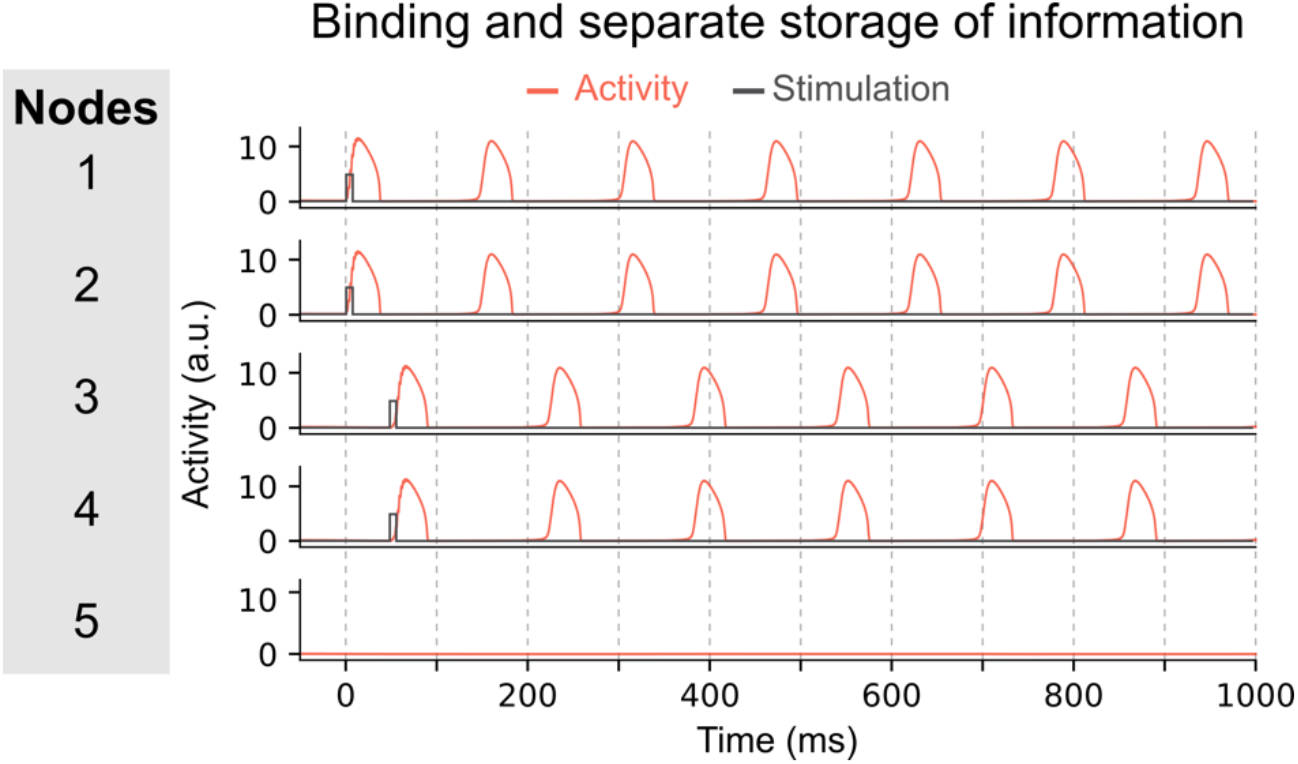
Maintenance of two pairs of items in the network. In this network, pairs of nodes are activated simultaneously, i.e. nodes 1 and 2 at time 0ms, and nodes 3 and 4 at time 50ms. Due to the inter-node coupling, each node of a pair will oscillate in synchrony and can thus represent a bound multi-element memory. The other pair will oscillate out-of-phase with the first pair allowing to store each binding separately and to permit read-out of each multi-element memory by downstream areas.

One of the parameters that determines the memory capacity, is the oscillatory frequency of the network, which itself is determined by the *τ* parameters. To illustrate the effect of frequency, we changed the temporal scale of the inhibitory component (*τ*_*v*_) of all nodes. In a first simulation (using *τ*_*v*_ = 32), each node oscillates at ~6 Hz once activated. The network can maintain up to 3 memories out-of-phase (i.e. their activity periodically peaks but never at the same time, allowing a downstream area to read-out each memory independently), see Figure 4a. In this network, the memory capacity is thus 3. In a second simulation we decreased the temporal scale of inhibitory components to obtain a network in which nodes oscillate at a slower frequency (*τ*_*v*_ = 20). In this network, activated nodes will oscillate at a faster frequency of ~10 Hz (see Figure 4b). When activating the first two nodes, they start oscillating out-of-phase, thereby maintaining 2 memories in the network. But when activating a third node, it will start competing with the first or second active node (depending on the exact timing of the stimulation of the third node). This competition will lead to one of three possible states: 1) the third node may not be able to sustain activation and this third memory will be lost; 2) the third node inhibits one of the other two nodes and the network will thus loose one of the previously stored memories; or 3) the third node synchronizes (or binds) with one of the activated nodes, thus creating a new bound memory. This last state is illustrated in Figure 4b. Each of these options shows that in this faster network, three distinct memories cannot be maintained concurrently, and therefore that a faster oscillatory frequency leads to a lower memory capacity.

**Figure 4.**
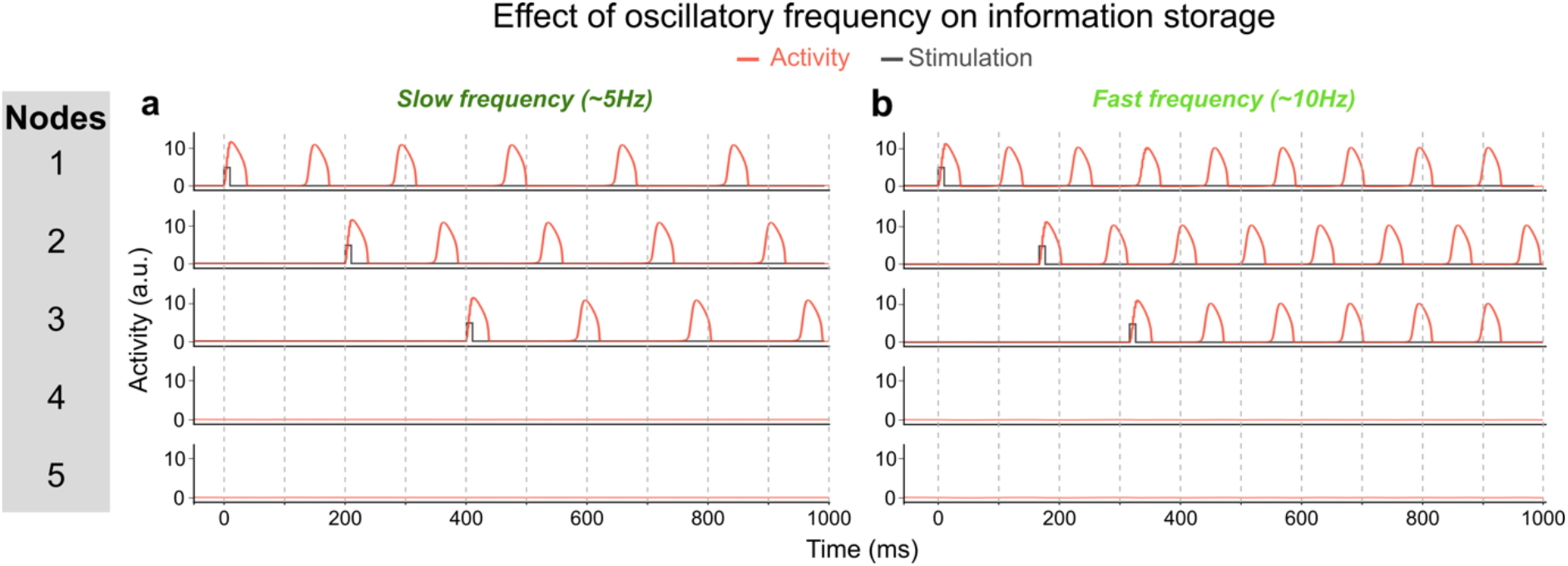
Effect of oscillatory frequency on maximum number of maintained items. **a.** In this network each activated node will oscillate at ~6 Hz (*τ*_*v*_ = 32). This allows to activate up to three units out-of-phase, i.e. three separately maintained memories. In this example, each node is activated sequentially and stays activated. **b.** In this network, the inhibitory time scale (*τ*_*v*_) has been decreased to 20 which increases the frequency of activated nodes to ~10 Hz. This prevents three nodes to be activated out-of-phase because of the short time interval between the activity peaks of activated nodes to “store” a third memory. In this example, node 3 is activated after nodes 1 and 2. Node 3 then shifts its phase and aligns with node 1, thereby losing the distinct information represented by each of the three nodes. This faster network thus has a lower memory capacity.

## General Discussion

To sum up, we have argued that synaptic and time-based binding have complementary advantages for the implementation of flexible cognition. Time-based binding can be quick and literally leaves no traces behind; but it leads to a natural processing (or resource) bound. Instead, synaptic binding is slower, prone to interference, but it does not suffer from this processing bound. We illustrated this theory with simulations of a recent oscillatory model. In the General Discussion, we relate our proposal to resource (or ‘bug’) theories, computational (or ‘feature’) theories, and to earlier oscillation theories. We end by pointing out some empirical predictions.

### Resource theories

In the current section, we discuss three influential theories on the nature of working memory constraints and resources, key data that earlier literature interpreted as supporting the respective theories, and how our own perspective accounts for those data.

A long-standing *slots theory* holds that working memory consists of a fixed number of slots (with the proposed number varying from 1 to 7) (Cowan, 2001; Miller, 1956; Zhang & Luck, 2008); one can consider slots as a discrete resource. A key behavioral signature interpreted as favoring slots theory, is the observation of a fixed precision for memoranda held in working memory beyond the slots upper bound. To be more specific, Zhang and Luck asked their subjects to retain from 1 up to 6 colors presented on different locations on the screen in working memory. In the test phase, they subsequently queried which colors were accurately remembered (subjects indicated the colors by clicking on a color wheel). Fitting a mixture model on the behavioral data in the test phase, the authors observed that the precision (inverse variance) of memory was statistically similar for 3 and for 6 items. They interpreted this as meaning that the number of available slots was equal to around 3. Instead, the precision in working memory did increase from 1 to 3; the authors interpreted this as meaning that more than 1 slot can be devoted to the same object (e.g., an object represented by two slots will be represented more precisely than one represented by just a single slot). However, fine-grained experimental paradigms with continuous manipulations of the relevant features (e.g., color or physical location), have since then demonstrated that there is no abrupt nonlinearity around 3 items (Ma et al., 2014). From the lowest set sizes on, when more items must be held in working memory, the representational precision of the remembered items gradually decreases. This data pattern was interpreted in terms of a *continuous resource theory* (often called resource theory in short), which holds there is a continuous but finite resource to be divided among the memoranda (Bays & Husain, 2008; Ma et al., 2014).

How to account for this data in a time-based binding perspective? We propose the following tentative theory. Suppose each neuron has a specific receptive field across some feature space (e.g., color space or Euclidean space). In the example, color- and location-sensitive neurons must bind to one another in order to represent the stimuli correctly. Suppose that there is a pool of neurons responding to specific colors and locations, in the color and Euclidean spaces, respectively. Suppose further that each neuron in one pool (e.g., responding to an active color) must be bound to at least one neuron from the other pool (responding to an active location) in order to influence downstream processing (i.e., be in working memory). Then, the precision will gradually decrease as more items must be retained: Indeed, more items retained means that less neuron pairs can be devoted to any specific item, given a finite period length of the oscillation. At first sight, it would seem that this theory predicts a hard bound at the maximal number of bindings that fit in a cycle, as in slots theory. However, given that several variables (including pool sizes, average receptive field, item location, period length, etc.), may vary from trial to trial, precision will also gradually decrease when additional items are in memory.

Another important behavioral signature is the occurrence of misbinding errors. This means that when (say) colors need to be remembered at specific locations, colors and locations may swap places in the participant’s memory. The existence of misbinding errors is naturally in line with a binding account; if two features (say, location and color) are incorrectly bound in memory, a binding error at behavioral level automatically follows.

The existence of misbinding errors was originally interpreted in terms of a third influential perspective on the nature of cognitive constraints on working memory, namely *interference theory* (Oberauer & Lin, 2017). Interference theory is in line with a long tradition of computational modeling via synaptic binding, specifically in neural networks. Interference theory holds that the postulation of a (discrete or continuous) resource is not required^2^. Cognitive processing in a neural network already leads to massive processing limitations due to (catastrophic) interference, and the latter is sufficient to explain the behavioral-level processing impairments that arise in tasks requiring the maintenance of several items at the same time.

At the risk of being overly reconciliatory, it’s worth pointing out that our time-based binding theory shares commonalities with each of the classical (discrete and continuous resources, interference) theories of working memory constraints. Like slots theory, it holds that just a fixed number of elements (here, bindings) can be maintained simultaneously. However, because the memoranda are bindings between neurons with variable parameters (cf. above), our perspective can predict, just like continuous resource theory, that there is no fixed bound at any number of items, and representational precision instead gradually decreases with more remembered items. Finally, binding elements together is rarely sufficient to solve actual tasks. The bindings must be read out by downstream task-specific processing modules. Such processing modules can most naturally be composed of standard neural networks, which implement synaptic binding, and are trained with gradient-based algorithms. The latter naturally also leads to similarities with interference theory.

### Computational-level theories

Generally speaking, computational-level theories consider that humans and other agents act in such a way as to achieve some goal (Lieder & Griffiths, 2020). As applied to working memory constraints, it holds that working memory may not be bound by the scarcity of a discrete or continuous resource, but that its boundedness is an optimal response to its environment. The early selection-for-action theory, for example, held that working memory must subserve action in the world (discussed in Hommel (2001)); and because action must be integrated (one cannot, for example, prepare a gratin dauphinois and play a video game at the same time), some environmental features must be selected, and others ignored.

More recently, Musslick and Cohen (2021) have combined computational-level and interference theory (see previous paragraph) to explain why cognitive control appears to be limited. Their starting point is the dilemma between learning and processing efficiency in standard neural networks: Efficient learning requires overlapping (shared) representations between tasks, but at the same time such overlap impairs multitasking (i.e., simultaneously performing two or more tasks). When tasks share input or output features, multitasking is almost impossible in such neural networks. Their simulations demonstrate that, in standard neural networks, basically just one task can be performed at any time. Thus, the optimal agent chooses to carry out just one task at a time. In their perspective, the bound is an optimal choice, given the computations one has to do and the architecture that is available for doing so. In principle, all tasks could be carried out at the same time; but because of the massive interference this would engender, an optimal agent chooses to limit the number of simultaneously processed items.

### Oscillation theories

The interest in neural oscillations dates back at least to the theoretical work of von der Malsburg (von der Malsburg & Singer, 1988). Around the same time, Gray, Singer, and colleagues detected that neural spiking in cells in the visual cortex phase-lock to the gamma rhythm (50-90 Hz); especially for features that are perceived as belonging to the same object (Gray & Singer, 1989). This led to the “binding by synchrony” and later the “communication through coherence” (CTC) theory. According to CTC, neurons in different brain areas can be bound together by firing in the same gamma phase (or more generally, by firing in a consistent and appropriate gamma phase difference; Fries (2015)). In particular, neurons in distant areas with a consistent gamma phase difference would share information more efficiently. For example, suppose the peak of the gamma wave is the phase where information can be most efficiently sent to other neural areas; if two neurons in distant areas always fire at this phase of the wave (i.e., the peak), this coincidence can be read out by downstream areas, and thus the two neurons are functionally (but not physically) bound. The binding of these different-area gamma waves could be orchestrated by a slower theta (4-7Hz) wave (Verguts, 2017; Voloh & Womelsdorf, 2016). Originally, this theory was proposed for the visual cortex, but has later been extended to cortical processing more generally. At this time, a massive amount of electrophysiological data supports (aspects of) the CTC theory (Womelsdorf et al., 2010), also in human cognitive control (Cavanagh & Frank, 2014), and in particular its relation with the slower theta wave (see also “Empirical predictions” section below).

In a second broad oscillation theory, Lisman and Jensen proposed that neural spikes that are locked to different gamma waves represent different pieces of information, where each gamma wave itself is locked to a different phase of the slower theta waves. This theory originates from findings observed in the hippocampus. In particular, the phenomenon of theta phase precession (O’Keefe & Recce, 1993) entails that as an animal proceeds in a cell’s preferred location, the cell’s spike firing time relative to theta phase moves earlier (precesses) in time. From this observation, it was proposed that the time of spike firing relative to theta phase is informative for downstream areas. This is what we will call the theta-phase binding (TPB) theory. Lisman and Jensen generalized this theta phase precession theory by proposing that items in working memory are stored by locking consecutive items in the list (each of them represented as neurons locked to gamma waves), to consecutive phases of theta (see also “Empirical predictions” section below).

Clearly, CTC and TPB theories have some commonalities; and they can be combined in the same framework, as was already demonstrated by McLelland and VanRullen (2016). Specifically, in a two-layer neural network model, they demonstrated that inhibition in the higher layer only, would cause patterns similar to what CTC would predict; whereas inhibition in both lower and higher layers, would instead cause patterns more similar to TPB. Also, our own time-based binding theory combines elements of CTC and PTB. Like CTC, it proposes that binding elements together is crucial for cross-area communication. Also, like CTC, it proposes that such binding is efficiently implemented via time. Like PTB, it holds that different packages of bindings can each be locked to a phase of a slower wave.

Finally, we note that, besides synapses and time, other binding schemes can be devised. For example, Akam and Kullmann (2010, 2014) proposed that also frequency could be used to bind elements together (as is also implemented in telecommunication systems). In general, any “labeling” of two or more elements would in principle be usable. Time and frequency happen to be the ones that are most naturally implemented via oscillations.

### Empirical predictions

Our perspective also leads to several empirical directions for future research. One interesting direction is to look at evidence for oscillations in behavioral measures. Recent literature has started to do just that by using dense temporal sampling paradigms in which the time interval between two events (e.g. a cue and a target) is varied across trials, allowing to estimate a time course of behavioral performance. For instance, Landau and Fries (2012) asked their subjects to pay attention to two horizontally lateralized gratings and notify the appearance of a brief contrast decrease. Spectral analysis (e.g. using Fast-Fourier Transform) of time-course of the accuracy data, obtained via a dense temporal sampling paradigm, revealed that attention fluctuated at theta frequency between the two gratings. Several other studies have replicated this finding (e.g. (Dugué et al., 2016; Fiebelkorn et al., 2013; Michel et al., 2021; Senoussi et al., 2019), and additionally expanded the study of oscillations in behavioral performance to the field of working memory (Peters et al., 2020; Pomper & Ansorge, 2021). This rapidly growing body of literature provides converging evidence that oscillatory processes are central to behavioral performance in a wide range of cognitive functions, in which they provide both a mechanism to sample or bind information, as well as a capacity limits of these functions.

In the field of working memory, predictions from the TPB theory have received support from several studies. According to the TPB theory, theta oscillations originating from medial temporal lobe and basal forebrain structures (e.g. hippocampus, septum) are hypothesized to support the maintenance of the ordinal information in an item sequence in working memory (Lisman & Jensen, 2013): the phase of theta oscillations structures the activation of distinct neural populations oscillating at gamma frequency, each representing an item of the maintained sequence. This theory thus predicts that a lower frequency of theta oscillations, leading to longer periods in which items could be nested, would lead to higher working memory capacity. Some studies have confirmed this prediction empirically by showing that higher working memory loads led to a reduction of theta frequency (Axmacher et al., 2010; Kosciessa et al., 2020). Moreover, a recent study causally tested this prediction using tACS (Wolinski et al., 2018) and showed that stimulating a fronto-parietal network at a slow theta frequency (i.e. 4Hz) led to higher working memory capacity than stimulating at a faster theta frequency (i.e. 7Hz). These studies confirm some predictions from the TPB theory as applied to working memory; and thus, they strengthen the view proposed in this article that oscillatory frequency modulates capacity limits in working memory, thereby constituting a factor limiting cognitive resources.

A related body of work has investigated the role of theta oscillations generated by the anterior cingulate cortex (ACC) in cognitive control processes (Cavanagh & Frank, 2014). Several studies have shown that these frontal theta oscillations are elicited when control is needed, i.e. during conflict or in preparation of a difficult task (Cavanagh & Frank, 2014), and allow to coordinate distant neural populations to create task-relevant functional networks through synchronization (Bressler et al., 1993; Canolty et al., 2006; Palva et al., 2005; Varela et al., 2001; Voloh & Womelsdorf, 2016). This theta-rhythmic process has been shown to support successful task performance (Voloh et al., 2015) and to support the instantiation of task rules (Womelsdorf et al., 2010). Critically, the frequency of these oscillations has recently been proposed to shift in response to task demands. Indeed, a recent study proposed that theta frequency balances reliable instantiation of task rules and the rapid gating of sensory and motor information relevant for the task at hand (Senoussi et al., 2020). They showed that this shift is observable both in oscillation of behavioral performance (using a dense behavioral sampling paradigm) and electrophysiological data, and that the magnitude of this shift correlates with inter-individual differences in task performance (Senoussi et al., 2020). Other studies have also reported the involvement of different low-frequency bands during top-down control processes, especially in hierarchical task implementation (Cooper et al., 2019; de Vries et al., 2020; Formica et al., 2021; Riddle et al., 2020). Together, these results open interesting avenue for future research on the conventional frequency limits of oscillations supporting cognitive control (usually attributed to the theta band) and more generally on the nature of the constraints controlling and limiting frequency shifts in neural oscillations. Future studies investigating the causes and consequences of frequency shifts in neural oscillations supporting cognitive control, for instance through neuromodulatory systems (Sara, 2015; Silvetti et al., 2018), will undoubtedly provide valuable insights on the neural bases of cognitive resources and their limitations.

### To conclude

We proposed that neural oscillations are both a solution to and a problem for flexible cognition. They are a solution because they allow items to be bound “on the spot”, leaving no synaptic traces that need to be erased afterwards, thus causing minimal interference (a notorious problem in standard artificial intelligence). They are also a problem because of the natural bound this system imposes; in this sense (only), the theory could be considered a resource theory.

## Supplementary information

The activity of each neuron of a neural triplet (u, v, and n) are defined by this system of differential equations:

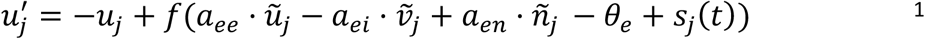

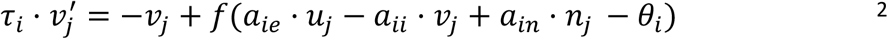

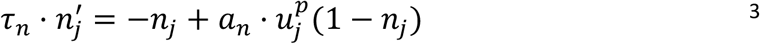

Where *u*_*j*_, *v*_*j*_ and *n*_*j*_ are the activity of the fast-excitatory, inhibitory and slow-excitatory components of node j, respectively. ***s***_***j***_(***t***) is the input signal (i.e. stimulation). Intratriplet coupling strengths are denoted by ***α*** parameters. Temporal constants are denoted by ***τ***, note that there is no temporal constant the fast-excitatory component (i.e. *τ*_*u*_ = 1). The parameters’ values are given in the table below:

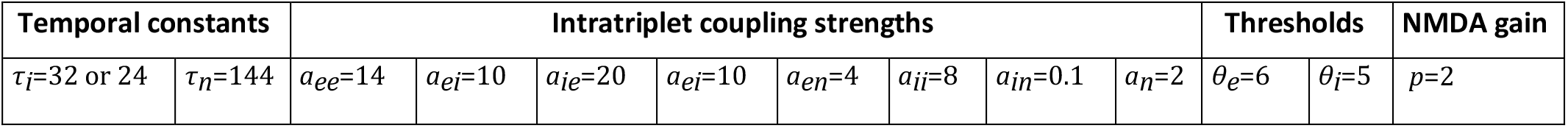

The function f(x) represents the firing rate (approximating a noisy-integrate-and-fire spiking neuron):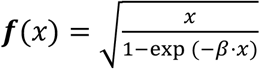.

The interaction between nodes (i.e. neural triplets) is defined by this equation:

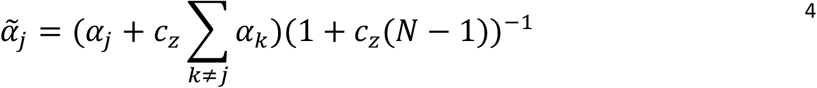

Where *α*_*j*_, denotes one of the three components of a triplet (i.e. *u*_*j*_, *v*_*j*_ or *n*_*j*_). Intertriplet coupling strengths are denoted by *C* parameters. The parameters’ values are given in the table below:

| Intertriplet coupling strengths |  | Number of nodes |
| --- | --- | --- |
| $c_e=0.001$ | $c_{ei}=0.03$ | $N=5$ |

## Footnotes

1 Without loss of generalization, we can assume *e* < *d*.

2 For completeness, it must also be mentioned that, in line with earlier work of Oberauer (2003), Oberauer & Lin (2017) also included a focus of attention in their model; an extra storage component that can hold just a single item. One could think of this storage component as a single slot (discrete resource).

